# Dynamics of Leukemic Blast and Immune Cell Populations in Acute Myeloid Leukemia

**DOI:** 10.64898/2026.03.21.713278

**Authors:** Sadiksha Adhikari, Philipp Sergeev, Nemo Ikonen, Minna Suvela, Heikki Kuusanmäki, Mika Kontro, Markus Vähä-Koskela, Caroline A. Heckman

## Abstract

Most patients with acute myeloid leukemia (AML) initially respond to standard chemotherapy. However, relapse and refractory disease remain common. The responses to targeted therapies are often transient and the efficacy of immunotherapy is limited. Although single-cell RNA sequencing (scRNA-seq) studies have provided insights into the cellular diversity and immune landscape of AML, many have primarily focused on limited, or newly diagnosed patient cohorts, leaving cellular dynamics across advanced disease incompletely defined.

Here, we profiled 72 samples from AML patients across different disease stages using scRNA-seq and compared these against healthy donor samples. We observed selective enrichment of immature progenitor populations, along with widespread upregulation of oxidative phosphorylation in AML. The immune microenvironment of AML was characterized by CD8+ effector memory T cell expansion with reduced IL2-STAT5 and increased mTORC1 pathways and exhaustion markers, suggesting a functional imbalance. Several AML-specific genes were identified providing potential therapeutic opportunities. Cell communication analysis revealed reduced HLA interactions in relapsed/refractory samples compared to diagnosis samples, suggesting impaired antigen presentation and defective T cell priming. Together, these results improve the understanding of cellular and immune changes in AML during disease progression and provide a basis for new therapeutic strategies.

## Introduction

Acute myeloid leukemia (AML) is a heterogeneous hematological malignancy of myeloid lineage with a complex cytogenetic and molecular landscape. The five-year survival rate is about 30% [1], with a severe prognosis for patients over 65 years of age [2]. Despite initial response to standard induction chemotherapy, the relapse rate is high and refractory disease is common [3]. Therapy resistant leukemic cells with self-renewal capacity [4], are major contributors to poor remission rate and efficacy of targeted therapies [5]. Increasing evidence suggests that AML progression and therapy resistance are shaped not only by intrinsic leukemic mutations but also by dynamic interactions between malignant progenitors and the bone marrow (BM) immune microenvironment [6,7]. Conventional AML treatments, including allogeneic hematopoietic stem cell transplantation (ASCT) and donor lymphocyte infusions, depend on the graft-versus-leukemia effects mediated by T-cells [8,9], highlighting the sensitivity of AML cells to the cytotoxicity of functional immune cells. While immunotherapy has revolutionized treatment of other cancers, its application in AML is still considered challenging, with limited efficacy and significant safety concerns. There is active research, promising early data, and some approved treatments, with ongoing efforts focused on making it safer and more effective [10]. Immunotherapy failure in AML has been linked to intricate heterogeneity of AML as well as low neoantigen burden, antigen overlap with healthy hematopoiesis, profound immune remodeling within the BM, T cells exhaustion, and defective antigen presentation. A promising approach for AML treatment is targeting immune checkpoints specifically expressed on leukemic stem cells (LSCs). Comprehensive studies are needed to improve our knowledge of the LSC-immune microenvironment interactions and provide critical insights for the development of more effective targeted and immunotherapy strategies.

While previous studies utilizing single-cell RNA sequencing (scRNA-seq) in AML have provided valuable insights into malignant cell heterogeneity and lineage organization, most have focused on newly diagnosed cases, limiting the representation of relapsed or refractory disease. This limitation has led to insufficient understanding of the extent of AML’s heterogeneity and molecular mechanisms related to therapy resistance. To address this gap, we characterize cellular composition, immune marker landscape and dynamics of leukemic cells and immune cells as well as their interactions with immune cells at single-cell resolution in a diverse cohort of AML samples at different disease stages. We also integrated single cell transcriptomes from healthy BM samples to unmask disease-specific cellular states and crosstalk.

## Materials and Methods

### Patients and clinical sample collection

We obtained 72 viably cryopreserved samples (71 BM and one peripheral blood (PB)) from 53 AML patients through the Finnish Hematology Biobank (www.fhrb.fi). The samples were collected with informed consent following ethical protocols approved by Helsinki University Hospital (study permits 239/13/03/00/2010,303/13/03/01/2011 and HUS/3235/2019). We also collected available associated clinical data including age, gender, subtype, treatment history, outcomes, survival, and karyotypes of the patients donating the samples. Mutation data was obtained with clinical data from the hospital and from sequencing data as mentioned previously [11].

### Single-cell RNA sequencing

Single cell gene expression profiles were generated using the 10x Genomics Chromium Single Cell 3’RNAseq platform. Library preparation and sequencing runs were conducted using the Chromium Next GEM Single Cell 3’ Gene Expression version 3.1 Dual Index chemistry. The sample libraries were sequenced on the Illumina NovaSeq 6000 system with read lengths of 28bp for Read 1, 10bp each for i7 and i5 index reads, and 90bp for Read 2. FASTQ files were produced with “cellranger mkfastq” from the 10x Genomics Cell Ranger v8.0.0 pipeline, and “cellranger count” was used to perform alignment, filtering, and UMI counting; mkfastq was run using the Illumina bcl2fastq v2.2.0. The scRNA-seq matrices were then analysed with the Seurat (v5) package in R. Dead cells with high mitochondrial gene expression were filtered. Doublets were removed with DoubletFinder (v 2.0). Cell annotation was performed by mapping the scRNA-seq data to a reference atlas in the BoneMarrowMap package [12], including batch correction with harmony within the package. Healthy donor BM scRNA-seq data [13], with unsorted BM cells similar to our cohort, including 25 samples from 20 donors, were integrated with our AML dataset. Progenitors were also called as blast cells in AML based on the expression of known AML markers [14]. We curated a list of immune marker genes (**Supplementary Table 1**) and explored their expression in different cellular compartments using DotPlot in Seurat. To explore if the genes in different groups are differentially expressed in different celltypes, we also compared their expression across sample subsets using a generalized linear mixed model (GLMM) with a Poisson distribution, accounting for random effects associated with individual samples. For multiple testing, post-hoc pairwise comparisons between conditions were performed using the estimated marginal means (EMMEANS) method.

### Differential gene expression analysis

Differential gene expression (DGE) analysis was conducted using the pseudobulk method with DESeq2 R package (v 1.48.1). A cutoff was established to ensure that a gene must be expressed in at least 5% of the total cells. Additionally, after DGE, filtering with log2 fold change cutoff of 0.25 was used. Genes expressed in fewer than 25% of cells in both groups were filtered. Genes with FDR corrected p-value less than 0.05 were considered significant. We performed gene set enrichment analysis (GSEA) using msigdbr (v.24.14.1) [15] and clusterProfiler (v 4.16.0) [16] using hallmark gene sets. Samples with ASCT in the treatment history and repeated samples from same patients were removed when comparing proportions of cells between different conditions. The ScTherapy tool was used to predict monotherapy drug response using differentially expressed genes results [17].

### Cell-cell communication analysis

For identifying cell-cell communication and differential interaction between the different cell states, CellChat (v 2.2.0) was used, with a minimum cutoff of 10 cells. We included only non-repeated samples to avoid pseudo-replication bias. Rare populations with very low representation in either the healthy or AML datasets (NK proliferating, T proliferating, stromal, and plasma cells; 4–66 cells across conditions) were excluded in differential interaction analysis to minimize noise during interaction inference.

### Scores

A Shannon diversity index was calculated with the vegan package in R using a proportion of each cell to measure intracellular diversity. Other scores were calculated using markers from the literature with UCell package [18] in R. A stemness score was generated using the CytoTRACE tool (v 1.1.0) [19].

### Statistical Analysis and visualization

For statistical comparisons, we employed the Wilcoxon test using the ggpubr package. Repeated measurements from the same patient within any group were avoided by including only diagnosis or the earliest sample after diagnosis Samples that received ASCT previously were removed when comparing immune cells.

### Flow cytometry

Viably cryopreserved MNCs from 148 patients with AML as well as 5 healthy donors were used for flow cytometry. After rapid thawing, the cells were pelleted by centrifugation (500g, 5 min) and the freezing medium removed. The cells were washed with complete medium (RPMI-1640 (ECB9006L, EuroClone), 10% FBS (A5256801, Gibco), 1% PenStrep (ECB3001D, EuroClone), 1% L-glutamine (ECB3000D, EuroClone)), and subsequently resuspended in completed medium supplemented with 12.5% conditioned medium (CM) from the HS-5 BM stromal cell line [20] , then incubated for 3h at 37°C and 5% CO_2_. To avoid cell clumping, 0.025% DENARASE (20804-1000k, c-LEcta) was added to the cell suspension. After incubation, viability of samples was measured, and samples were stained with respective antibodies and analysed by flow cytometry (FC). For staining, an antibody cocktail in staining buffer (SB, 1% FBS in PBS) was pre-plated using the acoustic liquid handler Echo 550 (Labcyte) and in 96-well V-bottom plates (Thermo Fisher Scientific). The antibody list can be found in **Supplementary Table 2**. Cells were centrifuged and resuspended in blocking cocktail (4% Human TruStain FcX (422301, BioLegend), 4% Tandem Signal Enchancer (130-099-887, Miltenyi Biotec), 1.33% IgG1 (400102, BioLegend), 1.33% IgG2 (400202, BioLegend) in SB) before mixing with antibodies. Cells in blocking cocktail mix were added to the pre-plated antibody mix (up to 250000 cells per well) in 1:1 ratio and incubated in the dark for 30 mins at room temperature on an orbital shaker (600 rpm). After incubation, the cells were washed with excess of SB, centrifuged and resuspended in SB for acquisition. Acquisition was done using the Novocyte Quanteon (Agilent) flow cytometer. Analysis of flow cytometry results was conducted using FlowJo v10.10 (BD Biosciences).

### Cell line data

Some genes of interest identified by scRNA-seq were explored using cell line data, especially to validate their expression in AML. For this, relevant expression data from DepMap portal [21] was extracted and explored in R.

## Results

### 1. Sample cohort and scRNA-seq data

We profiled a heterogeneous cohort of AML samples, including 22 Diagnosis (Dg), 2 remission (Rm), and 48 relapsed-refractory (RR) samples (**Table 1**) by scRNA-seq. Among these were 26 Dg-RR, 6 Dg-Rm-RR, and 4 RR-RR longitudinal sample sets. We integrated our AML scRNA-seq dataset with a publicly available healthy donor BM dataset. After preprocessing and filtering low quality, empty, dead cells as well as doublets, 312,302 high-quality cells, including 255,568 cells from AML and 56,734 cells from the healthy BM cohort were produced. The average number of cells for the AML cohort was 3,549 cells (558 - 7,750 cells) (**Supplementary Figure 1a**).

**Table 1.** Patient Characteristics.

| Sample | Stage | Subtype | Gender | ELN2022 | Genetic.characteristics |
| --- | --- | --- | --- | --- | --- |
| FIMM-AML-219_02 | Relapse | Not available | Male | Adverse | Chromosomal abnormalities not checked |
| FPM_AML_019_01 | Relapse | FAB M1 | Male | Adverse | Complex karyotype |
| FPM_AML_033_01 | Relapse | Not available | Female | Intermediate | Not available |
| FPM_AML_066_01 | Relapse | Not available | Male | Favorable | Not available |
| FPM_AML_070_01 | Relapse | FAB M2 | Male | Intermediate | 8 |
| FPM_AML_074_01 | Relapse | FAB M4 eos | Female | Favorable | inv(16)(p13.1q22) or t(16;16)(p13.1;q22), CBFB-MYH11 |
| FIMM-AML-210_02 | Relapse | Not available | Female | Not available | add(1q); del(20q) |
| FPM_AML_112_02 | Relapse | FAB M1 | Male | Adverse | Not detected |
| FPM_AML_116_00 | Diagnosis | Not available | Male | Not available | Not detected |
| FPM_AML_116_01 | Refractory | Not available | Male | Adverse | Not available |
| FPM_AML_117_01 | Diagnosis | FAB M1 | Female | Adverse | Not detected |
| FPM_AML_117_04 | Remission | FAB M1 | Female | Not available | Not available |
| FPM_AML_117_02 | Relapse | FAB M1 | Female | Not available | Not detected |
| FPM_AML_125_01 | Diagnosis | FAB M2 | Male | Adverse | del(5q) |
| FPM_AML_125_03 | Remission | FAB M2 | Male | Not available | Not available |
| FPM_AML_125_02 | Relapse | FAB M2 | Male | Not available | Not available |
| FPM_AML_128_01 | Diagnosis | FAB M2 | Male | Adverse | Not detected |
| FPM_AML_131_01 | Diagnosis | FAB M5 | Female | Intermediate | 21 |
| FPM_AML_131_02 | Relapse | FAB M5 | Female | Not available | Not checked |
| FPM_AML_133_01 | Diagnosis | Not available | Female | Favorable | Not detected |
| FPM_AML_141_01 | Diagnosis | FAB M1 | Female | Favorable | Not detected |
| FPM_AML_141_02 | Refractory | FAB M1 | Female | Favorable | Not checked |
| FPM_AML_142_01 | Diagnosis | FAB M1 | Female | Favorable | Not detected |
| FIMM-AML-211_02 | Relapse | FAB M1 | Male | Not available | Not available |
| FPM_AML_162_01 | tAML or sAML | FAB M2 | Male | Adverse | del(5q) |
| FPM_AML_162_02 | Refractory | FAB M2 | Male | Adverse | del(5q) |
| FIMM-AML-212_02 | Relapse | Not available | Female | Intermediate | Complex karyotype; t(8;10); t(1;10); abn(12) |
| FIMM-AML-213_02 | Refractory | FAB M5 | Male | Not available | Not checked |
| FIMM-AML-214_02 | Refractory | FAB M5 | Female | Not available | +8; +11 |
| FPM_AML_004_01 | Relapse | FAB M2 | Female | Favorable | Not detected |
| FPM_AML_006_02 | Relapse | FAB M2 | Male | Adverse | Not available |
| FIMM-AML-215_02 | Relapse | Not available | Male | Not available | +4; +22 |
| FPM_AML_015_05 | Diagnosis | FAB M5 | Male | Intermediate | t(1;5;14) |
| FPM_AML_015_02 | Relapse | FAB M5 | Male | Intermediate | t(1;5;14); +mar |
| FPM_AML_023_02 | Refractory | FAB M5 | Male | Intermediate | Not available |
| FPM_AML_035_01 | Diagnosis | FAB M1 | Female | Adverse | Complex karyotype |
| FPM_AML_035_02 | Refractory | FAB M1 | Female | Adverse | Complex karyotype |
| FPM_AML_038_01 | Refractory | FAB M2 | Male | Adverse | del(5q); +8; abn(8) |
| FPM_AML_040_02 | Relapse | FAB M5 | Male | Intermediate | t(5;11)(q35;p15.5) NUP98/NSD1 |
| FPM_AML_041_01 | Diagnosis | Not available | Male | Adverse | -7; t(3;3)(q21;26) |
| FPM_AML_043_02 | Refractory | FAB M2 | Female | Adverse | Complex karyotype |
| FIMM-AML-216_02 | Relapse | FAB M4 | Female | Not available | Not detected |
| FIMM-AML-217_02 | Relapse | FAB M2 | Male | Not available | 8 |
| FPM_AML_078_01 | Diagnosis | FAB M4 | Male | Adverse | Not detected |
| FPM_AML_078_02 | Refractory | FAB M4 | Male | Not available | Not detected |
| FIMM-AML-218_02 | Relapse | FAB M1 | Female | Not available | Not detected |
| FPM_AML_120_03 | Relapse | FAB M2 | Female | Intermediate | Not detected |
| FPM_AML_183_01 | Relapse | FAB M1 | Female | Intermediate | del(7p) |
| FPM_AML_183_02 | Relapse | FAB M1 | Female | Not available | Not available |
| FIMM-AML-200_01 | Diagnosis | Not available | Not avail | Not available | inv(3)(q21q56); -7 |
| FIMM-AML-200_02 | Relapse | Not available | Not avail | Not available | -7 |
| FIMM-AML-201_02 | Relapse | FAB M5 | Not avail | Not available | Not available |
| FIMM-AML-202_02 | Relapse | FAB M1 | Female | Not available | t(7;13) |
| FIMM-AML-203_01 | Diagnosis | Not available | Male | Not available | -3; -5; -13 |
| FIMM-AML-203_02 | Relapse | Not available | Male | Not available | -3; -5; -13 |
| FIMM-AML-204_02 | Diagnosis | Not available | Not avail | Not available | Not detected |
| FIMM-AML-205_02 | Relapse | FAB M1 | Not avail | Not available | del(6q) |
| FIMM-AML-206_02 | Relapse | FAB M5 | Female | Not available | Not detected |
| FIMM-AML-207_02 | Relapse | Not available | Not avail | Not available | tetraploidy |
| FIMM-AML-208_01 | Diagnosis | Not available | Not avail | Not available | Not available |
| FIMM-AML-208_02 | Relapse | Not available | Not avail | Not available | Not available |
| FIMM-AML-209_01 | Diagnosis | Not available | Not avail | Not available | Not available |
| FIMM-AML-209_02 | Relapse | Not available | Not avail | Not available | Not available |
| FIMM-AML-300_01 | Diagnosis | FAB M2 | Male | Adverse | Not available |
| FIMM-AML-300_02 | Relapse | FAB M2 | Male | Not available | Not available |
| FIMM-AML-302_02 | Relapse | FAB M0 | Male | Not available | Not available |
| FIMM-AML-302_03 | Refractory | FAB M0 | Male | Not available | Not available |
| FIMM-AML-303_01 | Diagnosis | FAB M4 | Female | Adverse | Not available |
| FIMM-AML-303_02 | Relapse | FAB M4 | Female | Not available | Not available |
| FIMM-AML-304_01 | Diagnosis | FAB M5 | Female | Favorable | Not available |
| FIMM-AML-305_02 | Relapse | FAB M5 | Female | Not available | Not available |
| FIMM-AML-306_01 | Diagnosis | FAB M5 | Female | Favorable | Not available |

### 2. AML displays extensive global remodelling of cellular composition and gene expression compared to bone marrow in healthy donors

As the basis for comparing AML and healthy donor samples, we mapped the integrated data to public reference map[13][15] and identified 25 major cell types (**Figure1a).** AML progenitor clusters were identified by expression of known AML markers (**Supplementary Figure 1b)**. The frequency of progenitors or blasts in AML samples correlated with the clinical blast percent (**Figure 1b)** and associated with FAB subtypes (**Supplementary Figure 1c)** When normalized to the total number of cells per donor, our results showed significantly reduced lymphoid populations and enrichment of progenitor populations in AML BM compared to healthy BM (**Supplementary Figure 2a)**. To determine lineage-specific changes, we analysed lymphoid and myeloid compartments separately. Compared to healthy donors, patients with AML showed an increase in NK cells, NK CD56 high subsets, and CD8 effector memory subsets (EM1 and EM2) (when normalized to total NK and T cells). In contrast, CD4 central memory, CD4 effector memory, CD4 naïve, and CD8 naïve populations were significantly reduced in AML (**Figure 1c).** Within the myeloid compartment, several progenitors remained significantly altered in AML samples (**Supplementary Figure 2b**). In addition, antigen-presenting cells, mature monocytes, and erythroid populations were reduced in the AML dataset.

**Figure 1.**
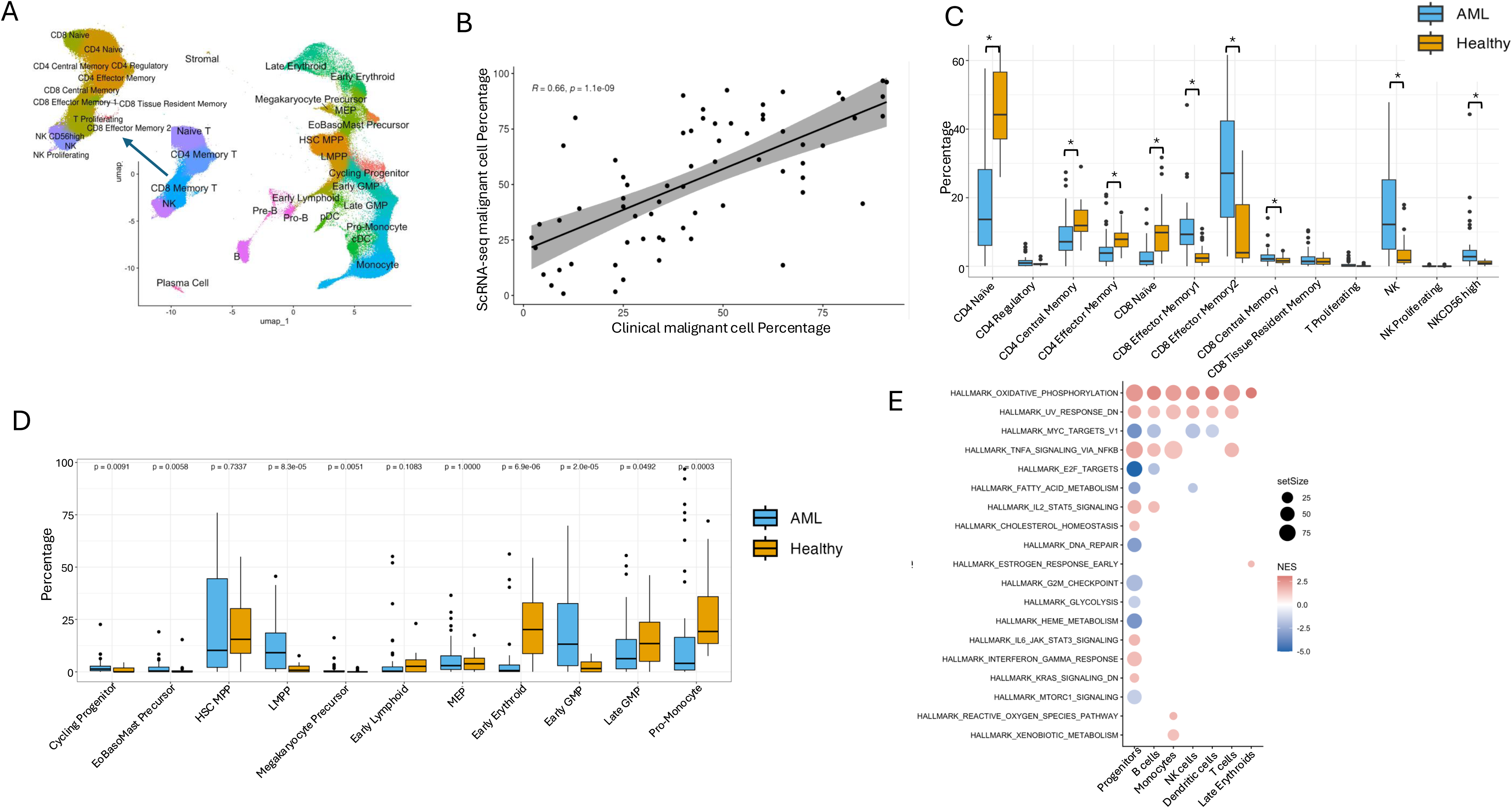
Overview of cellular compartments in AML and cellular dynamics in AML compared to healthy dataset. **a**. Cell clusters identified in AML. Annotation of lymphoid and myeloid cell subsets. **b**. Correlation between blast cells proportions identified by scRNA-seq with clinical blast percentage. **c**. Composition changes in T and NK cells. **d**. Composition changes in progenitor populations. **e**. Gene set enrichment analysis demonstrated pathways altered in AML compared to healthy BM. Positive NES refers to enriched pathways in AML.

Next, gene counts were normalized to total progenitor cells to identify AML specific enrichment in blast population. Progenitor population included Hematopoietic Stem Cell/ Multipotent Progenitor (HSC/MPP), Lympho-Myeloid Primed Progenitor (LMPP), cycling progenitors, Eosinophil-Basophil-Mast (EoBasoMast) precursors, megakaryocyte precursor, Megakaryocyte-Erythroid Progenitor (MEP), early erythroid, early and late Granulocyte-Monocyte Progenitor (GMP), pro-monocytes and early lymphoid cells. AML samples showed preferential enrichment of some immature progenitor states, including cycling progenitors, LMPP, and early GMP, whereas more differentiated progenitors including late GMP and pro-monocytes were reduced, when compared against healthy samples (**Figure 1d**). Differences in HSC/MPP, early lymphoid, and MEP were not significant. This suggests a selective increase of early progenitor states rather than uniform expansion across the hematopoietic hierarchy. Importantly, this compositional bias toward immature progenitor states was supported by functional transcriptome analyses: On one hand, LMPP, cycling progenitors, HSC/MPP, and early erythroid cells enriched in AML compared to healthy BM and exhibited high stemness scores calculated using by CytoTRACE (**Supplementary Figure 3a**), . On the other hand, functional state scoring further revealed heterogeneity within immature cell compartments, with HSC/MPP and LMPP populations predominantly labelled quiescent by the leukemia stem and progenitor cell (LSPC) score (**Supplementary Figure 3b**), whereas cycling progenitors displayed increased LSPC cycle scores (**Supplementary Figure 3c**). Together, these findings suggest that AML-associated enrichment of immature progenitor states includes both quiescent stem-like populations and actively proliferating malignant progenitors.

To provide a rough cell-lineage-enriched transcriptomic overview of AML *vs*. healthy datasets we next performed pseudobulk differential gene expression analysis between AML and healthy BM across major cellular compartment. The top upregulated genes in AML progenitors included *RACK1, GAS5, SNHG* family (*SNHG29, SNHG6, SNHG5*), immune and cytokine signalling genes (*TNFSF13, VSIR, NECTIN2, CD81,TNFRSF12A, TNFRSF10A, TNFRSF10B, TNFRSF10D, IFNAR1, IFNAR2, IL6ST, IL3RA*), as well as TGF-β pathway regulators (*TGFB1, TGFBR1, TGFBR2*) and *ADGRE* family genes (**Supplementary Table 3**). Pathway analysis showed that across nearly all AML cell types, oxidative phosphorylation was consistently upregulated (**Figure 1e**). TNFα signaling via NF|B and UV response pathways were also recurrently upregulated across multiple populations. In addition, progenitor cells showed suppression of pathways involving cell-cycle and metabolic pathways, including E2F targets, MYC targets, DNA repair, G2M checkpoint, fatty acid metabolism, glycolysis, heme metabolism, and mTORC1 signalling, along with enrichment of IL6–JAK–STAT3 and interferon gamma response.

To reveal possible actionable therapeutic differences between leukemic and healthy bone marrow cells, we utilized the scTherapy tool to predict potential monotherapies specific for AML progenitors. With DE genes in AML progenitors compared to the healthy dataset, 25 candidate drugs predicted high or moderate response, with alvocidib, ixazomib, camptothecin and docetaxel showing strongest predicted response across a mean of all samples **Supplementary Table 4)**. These drugs target mitotically active cells, consistent with AML progenitors upregulating several genes involved in cell cycle, DNA replication and protein synthesis (**Supplementary Table 3)**.

### 3. Cell-composition-based subgrouping of AML cases reveals gender-specific differences and differentiation-dependent drug vulnerabilities

As AML is heterogeneous, we wanted to assess if cellular composition could be used to subgroup the leukemic cases and if such subgroups correlate with clinical parameters or predicted drug responses. Myeloid lineage composition across samples in AML showed high inter-patient heterogeneity (**Figure 2a**). We identified four major cell phenotype classes: erythrocytic, monocytic, GMP/LMPP/early lymphoid-enriched and HSC/MPP-enriched phenotypes. The latter three classes could each be subclassified as mixed phenotype, depending on the level of cell types found across several of the main classes. While some samples were dominated by a single blast phenotype, others exhibited mixed compositions with malignant cells distributed across multiple cellular states, reflected by increased Shannon diversity indices. Interestingly, gender-associated differences were observed independent of a normalization strategy to total myeloid cells or blasts, with HSC/MPP-enriched states more frequent in males and late GMPs enriched in females (Wilcoxon test p < 0.05) . No significant age-associated differences in cellular composition were observed.

**Figure 2.**
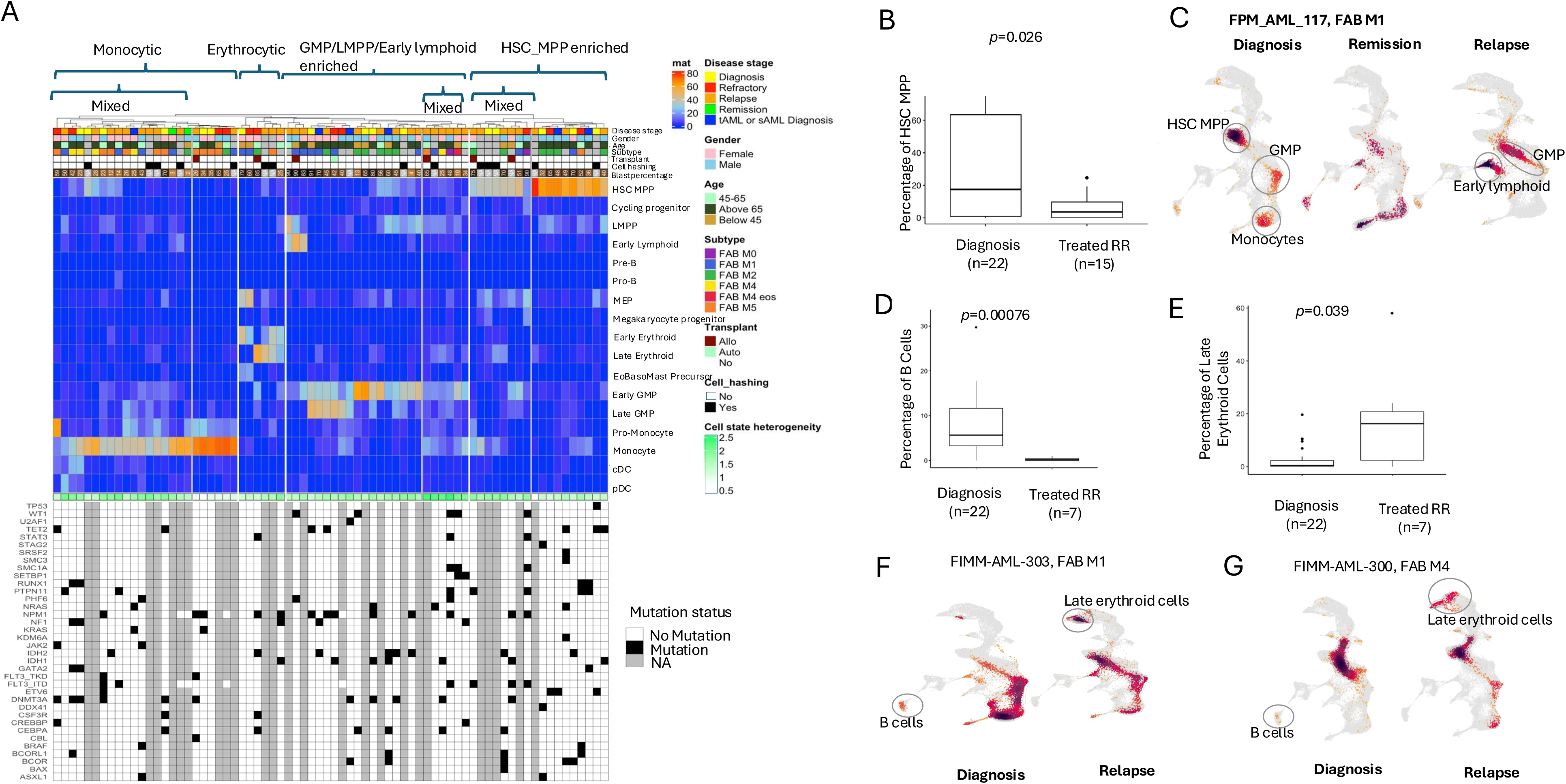
Cellular compostion in AML subgroups. **a**. Cellular heterogeneity among samples. Colors represent the proportion normalized to myeloid cells. Y-axis represents cell states and x-axis shows samples. Top annotation presents clinical data. Bottom annotation presents heterogeneity score calculated with Shannon diversity index and heatmap showing mutations in each sample. **B**. HSC MPP shows significant difference in at disease progression in patients treated with cytarabine based treatment. **C.** Cell population density changes in longitudinal samples from a patient treated with cytarabine-based regimen. **D-E**. Significantly different cell populations in samples from patients given venetoclax-based treatment compared to samples at diagnosis. **F-G.** Population density changes in longitudinal samples from patients treated with venetoclax-based regimen. P-value represents unpaired Wilcox test results.

When we compared predicted monotherapies using scTherapy in AML immature and mature cell types. We observed cell state-specific vulnerabilities distinguishing immature from mature blasts. Immature blasts were preferentially associated with sensitivity to epigenetic and proteosome inhibitors. In contrast, mature malignant cells showed kinase signalling dependencies. Several of these agents were predicted exhibit potentially toxicity (**Supplementary Table 4**).

### 4. Cellular composition of AML bone marrow is reshaped by chemotherapy and other treatments

Since our cohort included 26 Dg-RR, 6 Dg-Rm-RR, and 4 RR-RR longitudinal sample sets, with several treatments occurring between sampling (Table 1), we explored treatment-associated changes in cellular compositions during disease progression. We observed only minor changes associated with disease stage. Late erythroid cells were enriched in RR samples compared to Dg samples in myeloid lineage and among progenitors and lymphoid cells, MEPs and CD8 naïve cells were enriched respectively in Dg samples. (**Supplementary Figures 4 a-c**). In contrast, longitudinal sample analysis showed dynamic and heterogeneous AML evolution with case-specific directional shifts (**Supplementary Figure 4d**).

Next, we focused on specific treatment regimens. We restricted the analysis to Dg samples (n=22) and RR cases that had received cytarabine-based treatment (7+3 regimen) within 2 years prior to sampling and had not undergone ASCT (n=15). When normalized to blast cells, RR samples had significantly less HSC/MPP cells (**Figure 2b**) cells, while other stem-like blast cells (LMPP, early erythroid and cycling progenitor) were not significantly altered. Pseudobulk DGE analysis resulted only in differential expression of a few genes. While the reduction of HSC MPPs was observed in some longitudinal cases (**Figure 2c**), longitudinal samples showed heterogeneous changes reflecting the patient specific phenotypic heterogeneity and blast persistence **(Supplementary Figures 5a-e)**.

We also compared samples from the patients treated with 7+3 regimen by time to relapse (median >20 months). Blast cell gene expression of four long remission cases (range 37.87 to 84.93 months),when compared to five short remission cases (range 9.33 to 20.57 months), identified 50 genes (**Supplementary Table 5**). Short remission samples had upregulation of 15 ATP related genes, indicating enhanced mitochondrial function and increased ATP production, as well as upregulated *GPX1* expression, consistent with elevated oxidative stress adaptation. In contrast, *ANGPT1* and *EGR1* were downregulated, suggesting reduced growth signalling and angiogenesis.

Samples from venetoclax-treated patients (n=7) showed a higher proportion of late erythroid cells and a reduced proportion of B cells compared to samples taken at diagnosis (n=22) (**Figure 2d-e**). Two diagnosis-relapse longitudinal pairs, one with immature subtype and the other monocytic at diagnosis, both showed erythrocytic skewing at relapse (**Figure 2f-g**), while samples from relapsed patients treated with venetoclax showed loss of this population at the refractory stage (**Supplementary figure 5f**).

### 5. T cell remodelling reflects altered function of CD8 effector memory cells in AML

As mentioned earlier, comparison of the lymphoid cell proportion between AML and healthy donor samples revealed significant expansion of CD8 effector memory subsets (effector memory 1 and 2), alongside significant reductions in CD4⁺ central memory, CD4⁺ effector memory, CD4⁺ naïve, and CD8⁺ naïve populations.

To contextualize these shifts at the functional level, we first studied canonical memory differentiation within the AML microenvironment. We performed DGE and pathway analysis between memory *vs*. naïve cells separately for both CD4+ and CD8+ T cells, only in the AML dataset (**Supplementary Table 6**). In both analyses, we observed enrichment of IL2-STAT5 signalling along with other pathways (**Figure 3a-b**). However, when we performed DGE and pathway analysis directly contrasting expanded *vs*. reduced T-cell populations, the results showed distinct patterns (**Figure 3c; Supplementary Table 6**). Expanded CD8 EM populations had downregulated IL2-STAT5 signalling and MYC target pathway while enriching TNFα signalling via NF-κB and mTORC1 signalling compared to the reduced population. While these cells maintained increased expression of *STAT5A* and *STAT5B* expression, the functional pathway was compromised by significant downregulation of high affinity receptor *IL2RA*, and *MYC*. DGE results also showed upregulation of co-inhibitory markers (*CD160, TNFRSF9*), exhaustion markers (*TOX, LAG3* and others), cytotoxicity (*GNLY, FASLG* and others) and markers of NK cells (*NKG7, KLRD1, KLRG1*), pointing to NK skewing of CD8+ T cells. Interestingly, the checkpoint inhibitor CTLA-4 as well as its competitor *CD28* and marker of terminal differentiation *IL7R* were downregulated. Other interesting differences included downregulated *FLT3LG* (involved in T cell immunity and communication with dendritic cells) and *BCL2* (**Figure 3d**). Including NK cells in the expanded population (**Supplementary Table 6 )** preserved mTORC1 enrichment, but abolished IL2-STAT5 pathway downregulation (**Figure 3e**), indicating that IL2-STAT5 pathway suppression is specific to CD8 T cells. The IL2-STAT5 score, calculated using downregulated genes in this pathway, was low in all disease stages compared to the healthy dataset, while the exhaustion score was enriched in all disease stages (**Figure 3f-g**).

**Figure 3.**
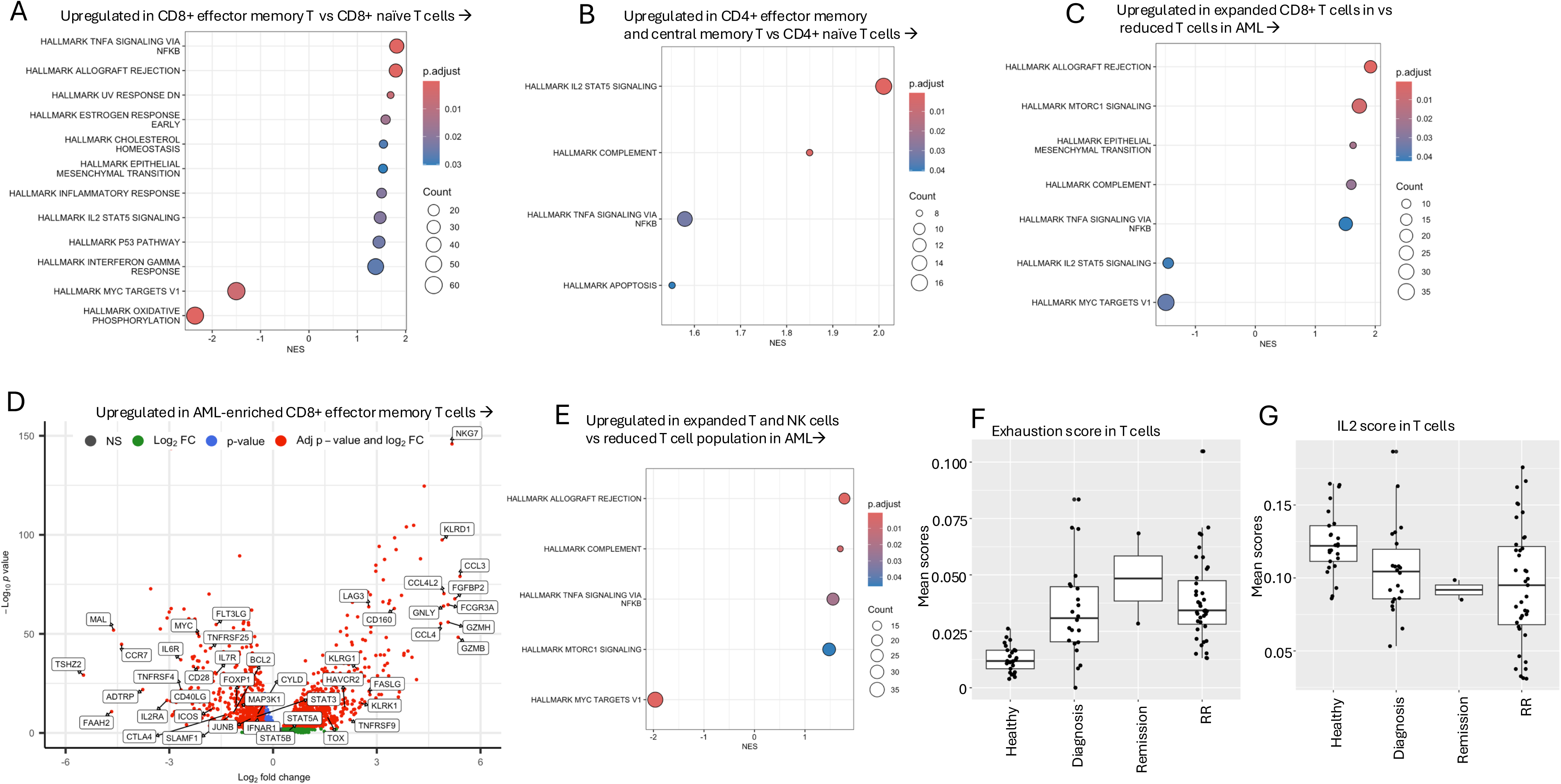
Immune compartment remodelling in AML. **A**. Hallmark pathways significantly different in CD8 effector memory T cells *vs*. CD8 naïve cells in AML. **B**. Hallmark pathways significantly different in CD4 effector and central memory T cells *vs*. CD4 naïve cells in AML. **C**. Hallmark pathways significantly different in T cells expanded in AML *vs*. T cells reduced in AML . **D**. Volcano plot showing top and bottom as well as selected significantly different genes in expanded *vs*. reduced T cells in AML. **E**. Hallmark pathways significantly different in combined expanded populations (NK and T cell subpopulations) *vs*. reduced T cells in AML. **F-g**. IL2-STAT5 and exhaustion scores in T cells in healthy and AML samples at different disease stages.

### 6. AML microenvironment presents altered leukemic–immune communication

Next, we performed cell-cell communication analysis in AML and healthy BM datasets to explore how cell-cell communication in the leukemic microenvironment diverges from the interactions observed in the healthy BM microenvironment. CellChat infers cell-cell communication using a curated signalling database by estimating ligand-receptor interaction probabilities and identifying significant interactions based on pathway level gene expression. Interaction strength is quantified as the aggregate of communication probability across the ligand-receptor interactions.

AML showed an increased number of interactions, but reduced interaction strength compared to healthy samples (**Figure 4a**). Differential interaction analysis showed that while cell-cell communication strength was mostly low in AML compared to healthy BM (**Figure 4b**), progenitors and monocytes exhibited the high outgoing interaction strength and clustered together (**Figure 4c**). These populations primarily signalled to NK cells, CD8 effector memory cells and NK CD56 high cells (**Figure 4c-d)**, whereas, CD8 naïve and central memory T cells had the lowest interaction strength. The upregulated signalling in AML included *MIF-CD74-CXCR4, CD55-ADGRE5*, *CLEC2B-KLRB1*, MHC-I-associated and other interactions **(Supplementary Figure 6a)**. These findings indicate distinct leukemic-immune communication compared to myeloid-immune interactions observed in healthy BM.

**Figure 4.**
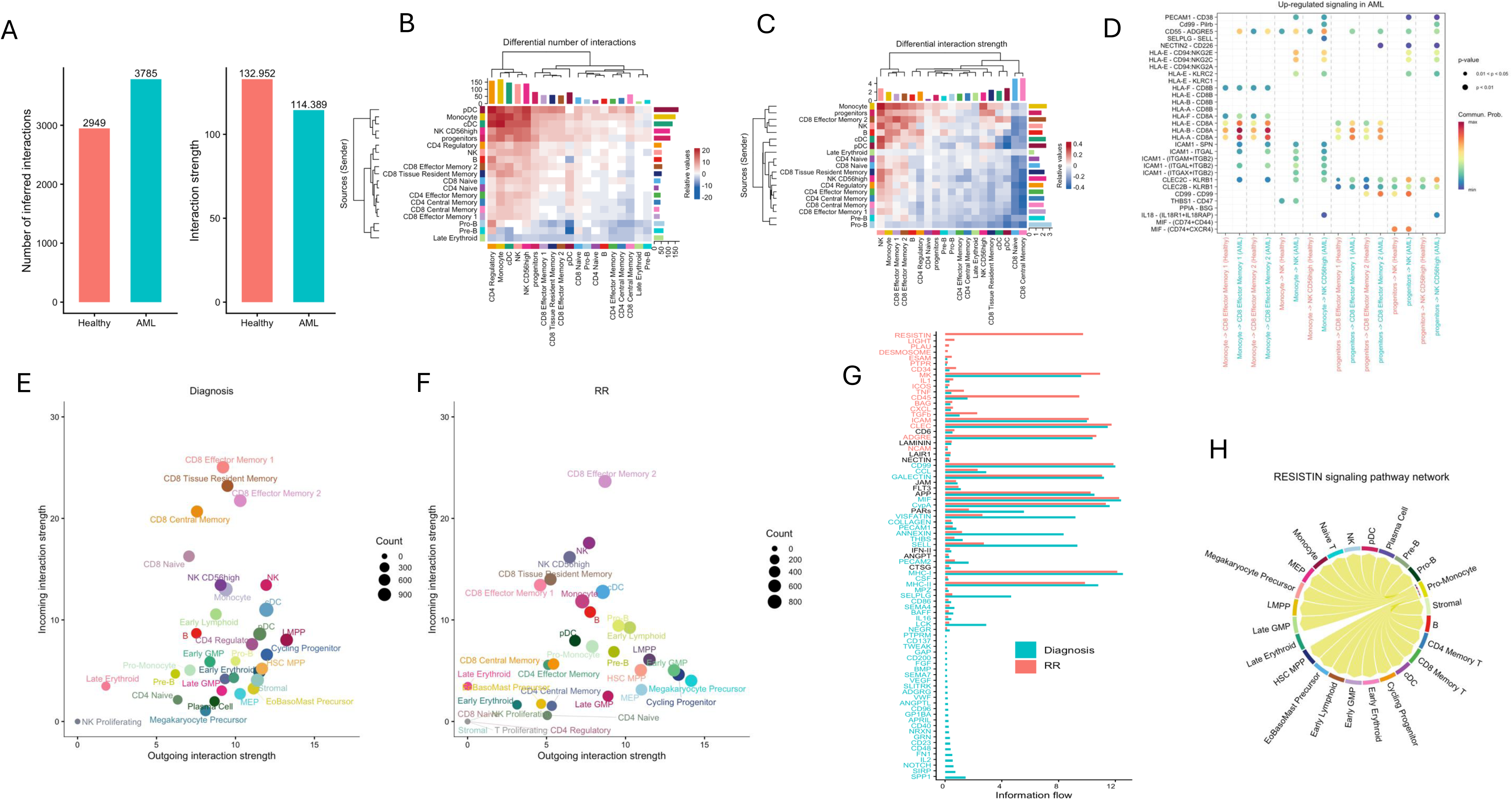
Cell-cell communication between different cell subsets. **A.** Number and strength of significant interactions in AML and healthy BM datasets. **B-C.** Differential number and strength of interactions in AML compared to healthy BM. Red represents enriched and blue represents downregulated cell-cell communication. In the heatmap, the top colored bar plot represents the sum incoming signalling to the cell types and the right colored bar plot represents the sum outgoing signalling from the cell types. In the colored bar within the heatmap, red and blue colors represent increased and decreased signalling respectively in the AML dataset compared to the healthy BM dataset. **D**. Heatmap showing signals from dominant sources to dominant targets in cell-cell communication that is enriched in AML. **E-F.** Cell types involved in incoming and outgoing integrations in diagnosis samples and RR samples. X-axis presents interaction strength (aggregated communication probability) of outgoing sources and y-axis presents that of incoming targets. **G.** Information flow (sum of interaction strength for each pathway) in diagnosis and RR samples. Blue represents interaction significant in diagnosis samples and red represents interactions significant in RR samples. **H.** RESISTIN interactions in AML RR samples.

Within the AML cohort, comparison of diagnostic and RR samples demonstrated fewer interactions and lower overall communication strength in RR compared to diagnostic samples (**Supplementary Figure 6b**). In both conditions, progenitor subpopulations remained the dominant signalling sources, and CD8 memory T and NK subpopulations remained dominant receivers (**Figure 4e-f**). Multiple signalling programs were shared between diagnostic and RR samples, including antigen presentation, pro-survival and inflammatory cytokines and adhesion/trafficking pathways (**Figure 4g)**. Notably RESISTIN signalling showed high interaction strength in RR samples driven by *RETN-CAP1* and *RETN-TLR4* interactions from pro-monocytes **(Figure 4h)**. In contrast, MHC interactions were reduced in RR, while TNF signalling was increased (**Supplementary Figure 6c-e**) along with *TGFB* and *LAIR1*. This indicates a shift from antigen presentation associated interactions towards immune regulatory signalling programs.

### 7. Expression of immune marker genes in AML is context specific

We next explored the expression of selected immune regulatory markers in myeloid and lymphoid compartments and performed GLMM for statistical significance, correcting for sample variability.

A broad shift in immune regulatory genes was observed in AML samples compared to the healthy cohort (**Figure 5a, Supplementary Figure 7a**). The most interesting results included expression of *VSIR, NECTIN2,* and *TNFSF13*, which were expressed only in AML samples and the latter two only in myeloid lineage, suggesting these to be potential targets for targeted therapy. Some markers such as *CD81* were upregulated in both myeloid and lymphoid populations in AML. In the lymphoid compartment, we observed upregulation of exhaustion-associated genes *TOX, TIGIT, LAG3* as well as co-stimulatory or activation-associated markers such as *ICOS* and *CD226* in naïve T cells, indicating coexistence of activating and inhibitory signals within the AML immune microenvironment. CD27 was downregulated in AML T and NK cells compared to healthy BM, consistent with altered T-cell maintenance signalling. Both *CTLA4* and *CD28* expression were higher in naïve and CD4 memory T cells in AML compared to healthy BM. The AML-enriched markers *VSIR, NECTIN2* and *TNFSF13* were expressed in both diagnosis and RR subgroups (**Supplementary Figure 7b**).

**Figure 5.**
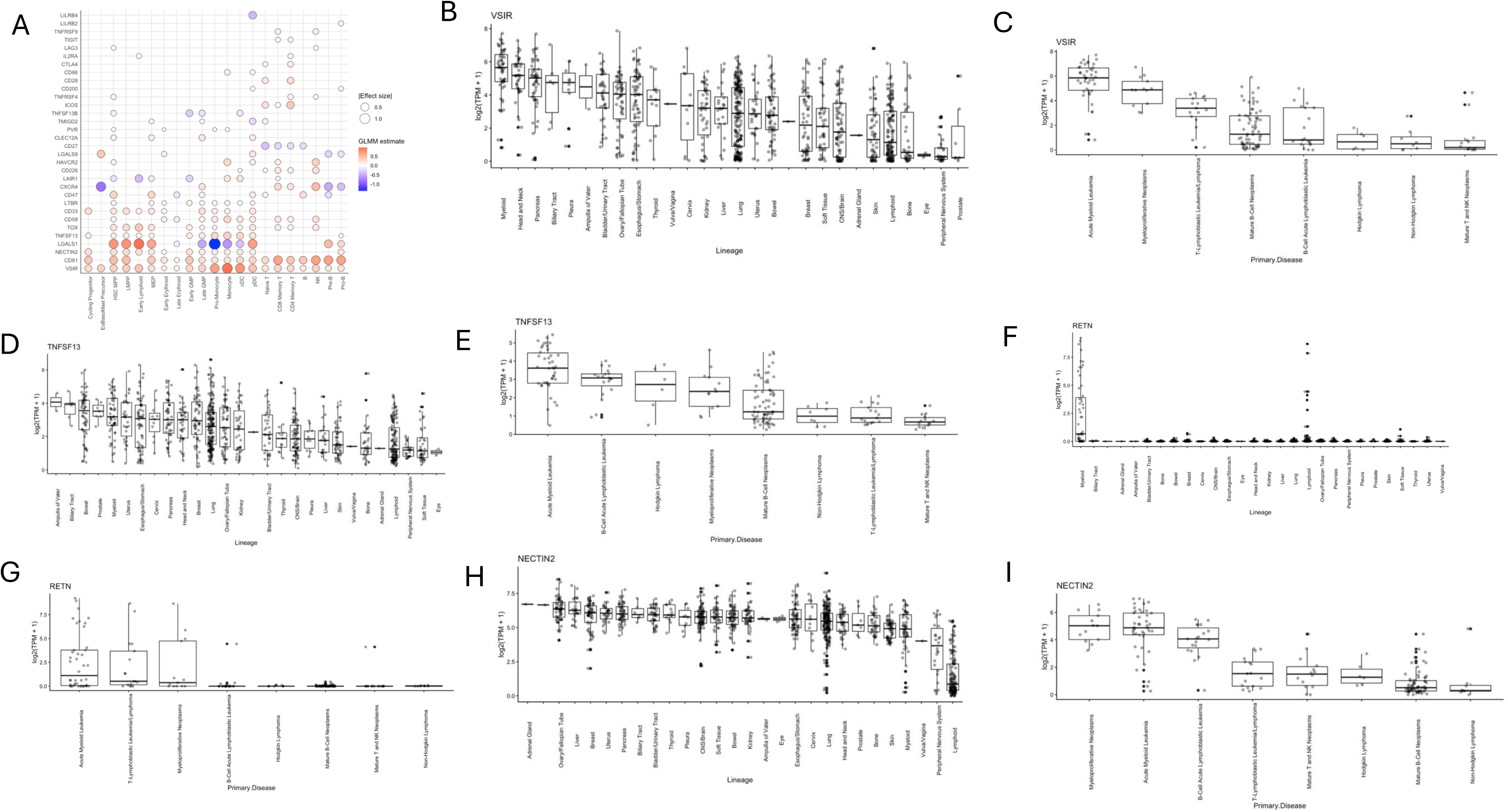
Immune marker genes with differential expression in AML cell subsets. **A.** Significant changes identified by GLMM analysis in AML compared to healthy datasets. Red represents increased estimate in AML and blue represents reduced estimate. Only results with p-value < 0.05 are plotted. Effect size is shown by dotplot size. **B-I** Expression of selected genes across different tissue types using cell line data from DepMap.

AML samples from cytarabine treated patients displayed reduction of few markers such as *CD200*, *CD58*, and *TOX* in multiple cell states in RR samples compared to diagnosis samples **(Supplementary figure 7c-e)**. Compared to diagnostic samples, samples from venetoclax-treated RR patients showed enrichment of multiple inhibitory and immune-regulatory markers across distinct cell populations including *LGALS1, VSIR, LAIR1, CD47, CD81, CD33* and *CD58* (**Supplementary figure 7f)**

We explored expression of selected AML and RR enriched markers in cell line data from DepMap portal. Interestingly, *VSIR/VISTA* had the highest expression in myeloid cells (**Figure 5b**). Among myeloid and lymphoid lineages, *VSIR* had the highest expression in AML, followed by myeloproliferative neoplasms (**Figure 5c**). *TNFSF13/APRIL* showed high expression in myeloid lineage cell subsets, while also being expressed in other lineages such as biliary tract, bowel, prostrate. Within myeloid and lymphoid lineages, *TNFSF13* also showed the highest enrichment expression in AML (**Figure 5d-e)**. *RETN* (RESISTIN) also emerged as an interesting potential target as it was found to be highly specific to the myeloid lineage (**Figure 5f-g**). *NECTIN2*, while showing high expression in multiple other lineages compared to the myeloid lineage, had highest expression in myeloproliferative neoplasms and AML (**Figure 5h-i**).

### 8. Orthogonal protein-level profiling of immune and progenitor cells in AML

We compared our results from scRNA-seq to multiparametric flow cytometry data across AML and healthy BM samples. Flow cytometry also presented broad differences in both progenitor and immune cell states in AML (**Supplementary Table 7**).

Comparing the cellular composition of AML, flow cytometry also suggested reduction in overall immune composition with significant loss of T cells with reduced lineage markers *CD3, CD4, CD27*, and activated memory cells marker *CD45RO* in AML compared to healthy BM. Further, significantly high PD1 expression was observed, validating the increase in inhibitory markers in AML. While results from flow cytometry demonstrated that Tregs were significantly higher in AML samples, in transcriptomics, we identified the trend but did not reach significance. Similar to transcriptomic data, erythroid progenitors were significantly reduced in AML, while early progenitors of myeloid and lymphoid lineage (CMP, CLP) were enriched. One notable difference was the reduced proportion of TIM3+ cells and CD4 naïve population in AML in the immunophenotyping data, whereas the transcriptomics data showed the opposite.

In RR samples from cytarabine treated patients, both transcriptomics and immunophenotyping data identified loss of HSCs compared to diagnostic samples. Both datasets showed reduction of exhaustion markers in the T cell compartment, with flow cytometry data showing reduced TIGIT and transcriptomics data showing reduced *TOX* in CD8 populations in RR samples. Some variations were noticed. Notably in flow cytometry data, TIM3 was significantly high in RR samples from treated patients, but such changes were not identified in the scRNA-seq results. With regards to venetoclax treatment, both datasets suggested significant increase in exhaustion marker expression; particularly TIM3 in flow cytometry data and *TIM3 and VSIR* in transcriptomics data. Similar to scRNA-seq, flow cytometry results showed CD47 was enriched in samples from venetoclax treated patients, suggesting blast escape from immune recognition.

### 9. Exploratory analysis identified associations with recurrent AML mutations

Next, we explored the association of AML mutations with cellular composition changes and immune marker expression. We observed several mutation-associated changes.

In *DNMT3A*-mutated AML (n=8), scRNA-seq analysis showed increased proportions of pro-monocytes compared to wild-type samples (n=24) (**Supplementary Figure 8a-c**). In the immune compartment, CD4 naïve T cells and CD8 effector memory 1 T cells were enriched in mutated samples, which was also validated by flow cytometry. At the immune-regulatory level, *DNMT3A* mutations were associated with increased *TIM3, CXCR4, CD58*, and *CD200* expression in specific progenitor and lymphoid subsets, elevated *TOX* in pro-B cells, and higher *CLEC12A* in pDCs, while *LILRB2* and *CD47* were reduced in monocyte populations, with some results such as *LILRB2* also validated by flow cytometry. *IDH1* mutations (n=6) were associated with reduced monocytes and cDCs compared to wild-type (n=26) (**Supplementary Figure 8d-e)**. At the immune-marker level, *IDH1* mutation was associated with increased *CD58* and *HAVCR2* (*TIM-3*), was also observed in the immunophenotyping results, in addition to *TOX* and *KLRG1* in several cell states. Conversely, *LILRB4, CXCR4, CD86* and *CD33* were reduced in multiple cell types. *IDH2*-mutated samples did not show significant compositional changes. Immune-marker analysis revealed lower *LGALS1, CD33*, and *CD200* expression in multiple cell types, with increased *HAVCR2* in early erythroid cells, *ICOS* in naïve T cells, and *TNFRSF4* and *PVR* in some cell states. Both *IDH1* and *IDH2* mutations were associated with reduced *LGALS9* expression in monocytes.

*CEBPA*-mutated samples (n=4) showed increased MEP abundance compared with wild-type (n=27) in scRNA-seq (**Supplementary Figure 8 f)**, and flow cytometry indicated enrichment of CD8 T and CD4 central memory T cells. *NRAS*-mutated samples (n=3) showed reduced NK cells, CD4 regulatory T cells, and CD8 central memory T cells compared with wild-type (n=28). *PTPN11*-mutated samples (n=3) showed enrichment of early lymphoid blast populations and NK cells (wild type n=28) (**Supplementary Figure 8g-h**). *NPM1*-mutated samples (n=8) showed reduced cDC proportions, compared to wild-type (n=23), although proportions were low in both groups (**Supplementary Figure 8i)**. Immune-marker analysis revealed higher *CD33, LAIR1, TNFSF13B (BAFF)* and *CD70* expression across different cell types and reduced *TMIGD2* and *HAVCR2* expression in mutated samples. NF1-mutated samples showed increased *CXCR4*, *CD47*, and *LAIR1* expression across multiple myeloid states, along with increased *CD27* in CD8 T cells and *NECTIN2* in pro-B cells. Reduced *LAIR1* in early erythroid cells and reduced *CD47* in pro-B cells were also observed. *FLT3*-ITD–mutated samples demonstrated enrichment of megakaryocyte precursors, although at low absolute frequencies (**Supplementary Figure 8j)**. Immune-marker analysis showed increased *LGALS1, KLRG1*, and *IL2RA* (*CD25*) expression in mutated samples, while *CXCR4* expression was higher in wild-type. No major compositional differences were observed in *BCOR*-mutated AML. However, immune-marker analysis showed altered *CXCR4* distribution across T-cell subsets, increased *LGALS1* expression in B cells in and reduced *LILRB2/4, TNFSF13B,* and *CD86* expression in myeloid cells of mutated samples. A comprehensive list of associations between immune marker expression and mutations in AML based on scRNA-seq data is presented in **Supplementary Table 8.**

Flow cytometry showed additional mutation-associated differences in cellular composition for multiple mutations. *FLT3* mutated samples showed reduced CLP, MEP, and T-cell populations except CD8 T cells. In *FLT3*-ITD mutated samples GMPs were enriched. *IDH2* mutated samples had reduced T cells, increased LSC, *LMPP* and Treg, *IDH1* had reduced LMPP, CD8 naïve and increased CD8 central memory cells. *NPM1* mutations were associated with increased CLP, GMP, CD4+ TE, and CD8 cells as well as reduced HSC, MPP, MEP and naïve T cells. *RUNX1* mutated samples had enriched HSC and MPP. *GATA2* mutated samples showed enriched myeloid progenitors, and CD8 cells. Flow cytometry further demonstrated context dependent changes in immune marker expression. For example, CD47 expression was increased in *IDH2* mutated cases, and reduced in *FLT3* mutated cases. CD58 was reduced in *RUNX1* mutated cases, CD74 increased in *NPM1* mutated and *IDH2* mutated cases, but decreased in *IDH1* mutated cases. LILRB2 *was* enriched in *IDH1, IDH2, CEPBA* mutated samples but reduced in *RUNX1* and *NF1* mutated cases. Complete immunophenotyping results are summarized in **Supplementary Table 9**.

## Discussion

In this study, we performed integrated single-cell transcriptomic profiling and proteomic analysis of a large and clinically diverse AML cohort to explore cellular composition and diversity, cell specific gene expression changes, as well as immune-leukemia communication and immune marker changes. Our data suggest that AML reshapes the hematopoietic hierarchy and coordinates metabolic and immune rewiring across cellular compartments. While we also observed increased heterogeneity in malignant cell populations in AML samples [12,22–25], we observed non-uniform composition changes across the hematological hierarchy with selective enrichment of early progenitors such as LMPP and cycling progenitors and reduction in more differentiated progenitors in AML compared to the healthy dataset. Across multiple cellular compartments, we observed enrichment of oxidative phosphorylation associated pathway enrichment in AML.

In our cell-cell communication analysis, AML enriched interactions included activation (*NECTIN2-CD226* and *HLA-E-CD94/NKG2C*) and inhibitory (CLEC2B-KLRB1) and cell survival (MIF-CD74-CXCR4) signalling, highlighting complex and functionally diverse leukemic-immune signaling landscape. *ADGRE* signalling was significantly high in AML. Targeting ADGRE family genes have been suggested to remove off target toxicity as combination treatment with CAR-T therapy [26,27]. Despite quantitative differences in interaction number and strength between cell-cell communication signalling in diagnosis and RR samples, many of the signalling pathways were shared, indicating preservation of most communication network across disease stages. A shift in signalling intensity and cellular contributors was observed in RR stage. For example, both MHC-associated interactions were reduced in the RR compared to diagnosis group, suggesting impaired antigen presentation and T-cell priming. Defective antigen presentation prevents T cells from recognizing and attacking tumor cells, and is a primary mechanism of resistance to immune checkpoint inhibitor therapies despite the removal of inhibitory signals [28]. RESISTIN signalling was enriched in RR samples. Higher *RETN* expression has been reported previously in pediatric acute lymphoblastic leukemia relapse cases compared to initial diagnosis or healthy controls [29]. Interestingly DepMap data showed high *RETN* expression in myeloid lineage cell lines, especially AML, suggesting this as a potential therapeutic target.

We also explored functional transcriptomic programs within T-cell subsets expanded in AML. By separating lineage-specific differentiation from AML-associated selection, our analyses reveal that preferentially expanded T cells in AML are not defined simply by memory status or activation, but by distinct pathways. T cells that accumulate across the AML niche exhibited enrichment of mTORC1 pathway with reduced MYC signalling and IL-2–STAT5–mediated pathways compared to T cells that were reduced in AML. Enrichment of the mTORC1 pathway is representative of high metabolic activity and plays a large role in T cell activation. However, prolonged, activated mTORC1 can lead to functional defects that prevent these cells from reaching and killing leukemic blasts, while lowering mTORC1 activity has shown promising effects in CAR-T therapy for AML [30]. It’s also been shown that while mTORC signalling may still be active upstream, the downstream signalling and cellular responses necessary for T cell proliferation and function can be compromised due to reduced MYC activity, leading to impaired metabolic reprogramming and possibly T cell exhaustion [31]. In line with this, AML-enriched T cells showed enrichment in exhaustion markers such as *TOX, LAG3 and TIGIT,* along with markers of NK-cells (*NKG7, KLRD1, KLRG1*) and downregulation of STAT3. A NK-like signature of T cells has been discussed before in AML to promote T cell dysfunction and persistence, which is different from solid tumor-associated canonical exhaustion programs [42]. Reduced *STAT3* also suggests compromised effector function and terminal differentiation as *STAT3* has been shown to promote the effector function of CD8+ T cells and has a role in terminally differentiated CD8+ T cell development [32]. Decreased IL-2 production and STAT5 phosphorylation level have been discussed in depleted CD4 T cells [33]. While IL2 monotherapy has limited efficacy and high toxicity, some research suggests modifying the structure or presentation of IL-2 can reduce toxicity and lead to effective anti-tumor responses in synergy with checkpoint blockade [34]. Recently, it has appeared as a promising therapy in low dose and combination for AML subgroups [34]. Together our findings suggest deregulated and functionally impaired T cells in AML with potential therapeutic relevance.

We also accessed context-dependent immune regulatory genes expression in AML. *VISTA* [35] and *CD81* [36], targets of clinical and preclinical studies, were present in both immune and myeloid compartments, with higher expression in AML compared to healthy BM. While CD81 expression has been discussed in blast cells [36], to our knowledge it has not been equally discussed in T and NK cells. *TNFSF13 (APRIL)* and *NECTIN2*, enriched in AML, were myeloid-lineage specific and may serve as potential clinically significant diagnostic markers to detect leukemia or potentially present as a target with low toxicity. DepMap cell line data suggested these genes are highly specific to AML and therefore requires further exploration. *TNFSF13* has been identified as a positive regulator of AML initiating cells in a murine-based study [37], and found to be involved in immunosuppression in gliomas [38]. It’s receptor, BCMA, is the target of several approved immunotherapies and has recently gained attention as a promising novel antigen in AML [39]. Inhibition of *TNFSF13* signaling has been reported to significant increase the apoptotic cell death induced by camptothecin, independently of the subtype of the leukemia samples [40]. Interestingly, in our scTherapy drug prediction results, campothecin was one of the drugs predicted to have high to moderate response on AML blast cells. While our top drug predictions based on scRNAseq revealed mostly antimitotics targeting CDK:s, proteasome, topoisomerases and microtubules (Supplementary Table 4), which are also toxic to healthy cells, it could still be possible to create new cocktails of these agents with an overall greater therapeutic index than achievable with current induction chemotherapy. Such a hypothesis would require extensive experimental exploration beyond the scope of the current report. Other genes such as *LGALS1* were found to be expressed in AML progenitors. *LGALS1* gene expression has been identified as an independent unfavorable prognostic factor, and its inhibitor OTX008 has shown significant therapeutic effects [41].

Therapy and mutation-associated analyses on cellular composition and immune related gene expression further demonstrated that ecosystem remodelling is dynamic and context-dependent. *CD200* has been discussed only in *NPM1* mutated samples but not associated with cytarabine-based treatment to our knowledge [42] . Consistent to our results, increased *CD33* expression has been well-documented in the context of *NPM1* mutations [43]. We observed that the patient’s phenotype may change, with cytarabine-based treatment mostly targeting stem-like cells as shown by loss of HSCs. We observed that samples from patients with short response duration to cytarabine-based treatment had upregulation in 15 ATP genes. Extracellular ATP along with CD39-P2RY13-cAMP-OxPHOS axis have been discussed as key regulators of cytarabine resistance [53]. Patients treated with venetoclax-based regimen showed increased erythroid differentiation at disease progression. Erythroid cells highly express and depend on BCL-XL for survival; as venetoclax selectively inhibits BCL2, AML cells with erythroid or megakaryocytic differentiation exhibit reduced sensitivity to venetoclax [44]. Earlier investigations have also demonstrated erythroid differentiation could be promoted by *CEPBA* mutations [45]. In our study, *CEPBA* mutations were also linked to higher proportions of MEPs in our study. Given the limited sample sizes for individual mutations, with regards to mutation and immune markers our results presented associative trends rather than definitive mutation-specific phenotypes.

Although these analyses were exploratory and limited by subgroup size, the observed shifts in progenitor composition after cytarabine-based treatment, erythroid skewing after venetoclax-based treatment, and immune marker expression support the concept that distinct selective pressures shape AML cellular ecosystems in a non-uniform manner. Collectively, our data refine the structure and organization of AML-associated dynamics across cell states and clinical contexts, suggest potential therapeutic targets, and provide a framework for future studies investigating how malignant progenitors and immune cells co-evolve within the leukemic niche. Future longitudinal and functional studies will be required to clarify the clinical implications of these cellular and molecular changes in AML.

## Acknowledgement

We are thankful to the patients who donated samples for the study and Finnish Hematology Registry and Biobank for providing the samples. We acknowledge the assistance and support of the FIMM Sequencing Unit and Single-Cell Analytics Unit, particularly for their help with sequencing. The study was supported by grants from the Sigrid Jusélius Foundation, Cancer Foundation Finland, and the Research Council of Finland (grant no. 1357686 and 1320185).

