## Supplementary Figures for "Dynamics of Leukemic Blast and Immune Cell Populations in Acute Myeloid Leukemia"

#### Supplementary figure 1. Cellular composition in AML

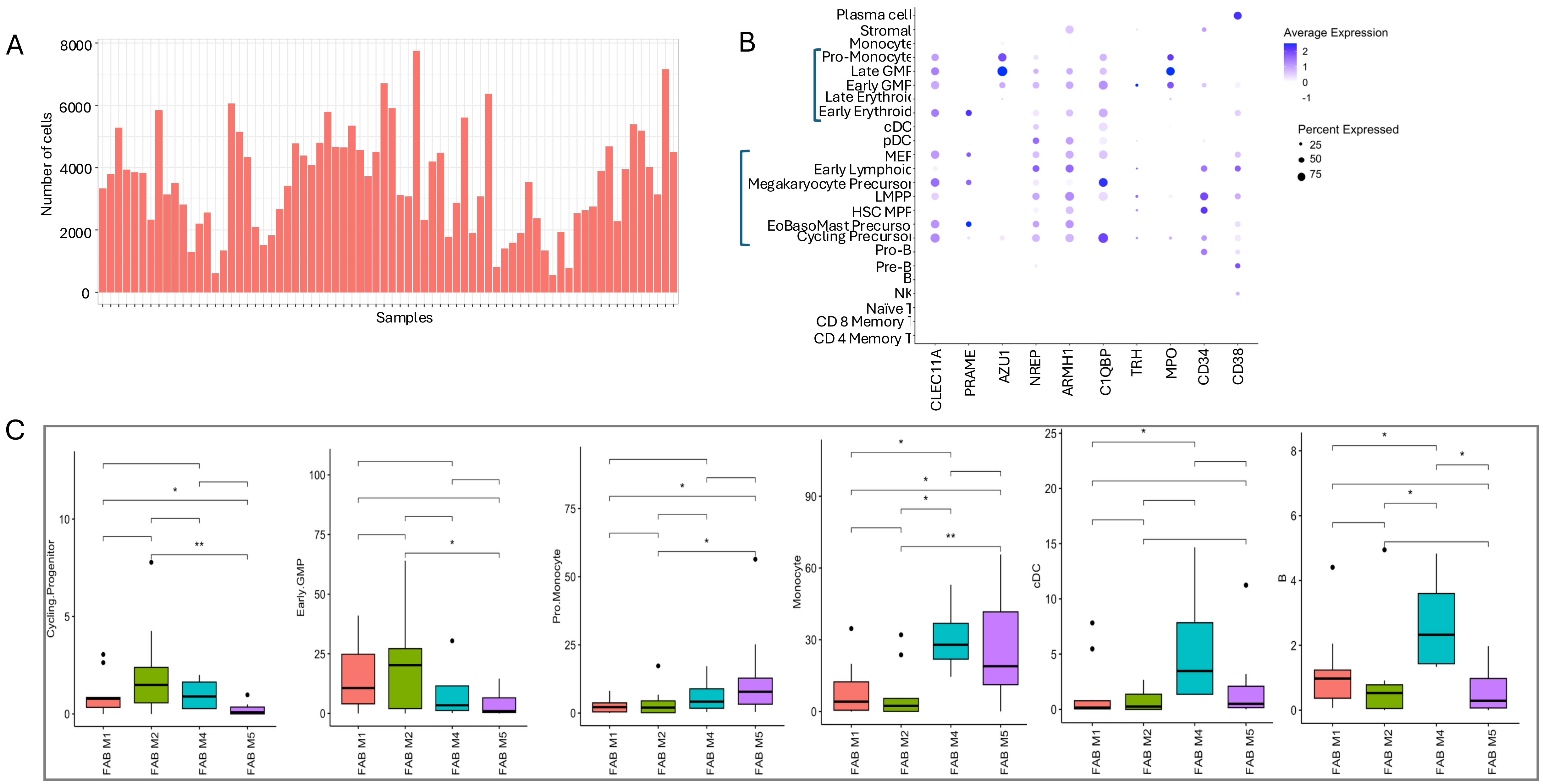

**Supplementary figure 2. Composition shift in AML compared to healthy BM samples.**

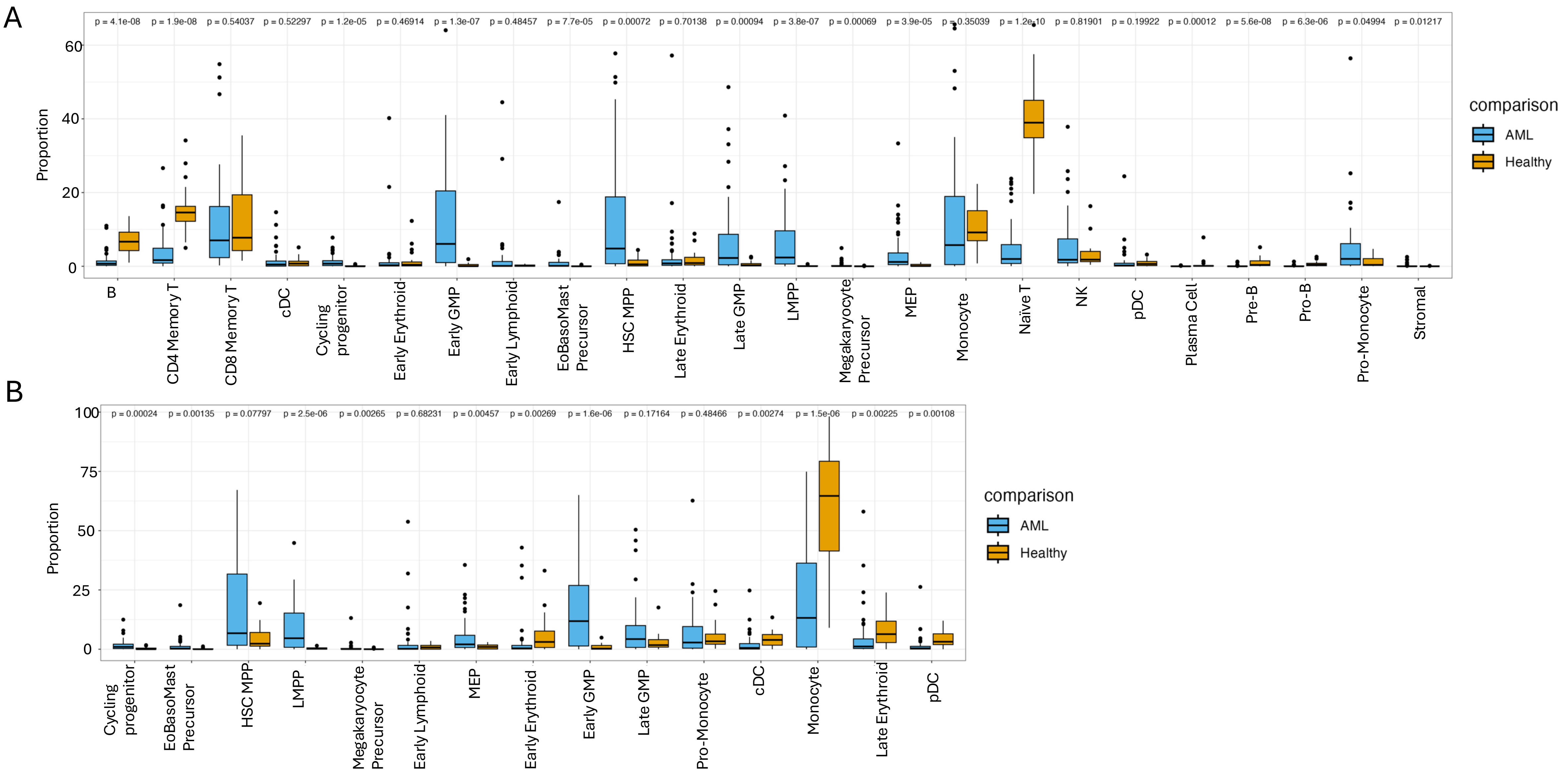

**Supplementary figure 3. Functional states in AML myeloid cell populations**

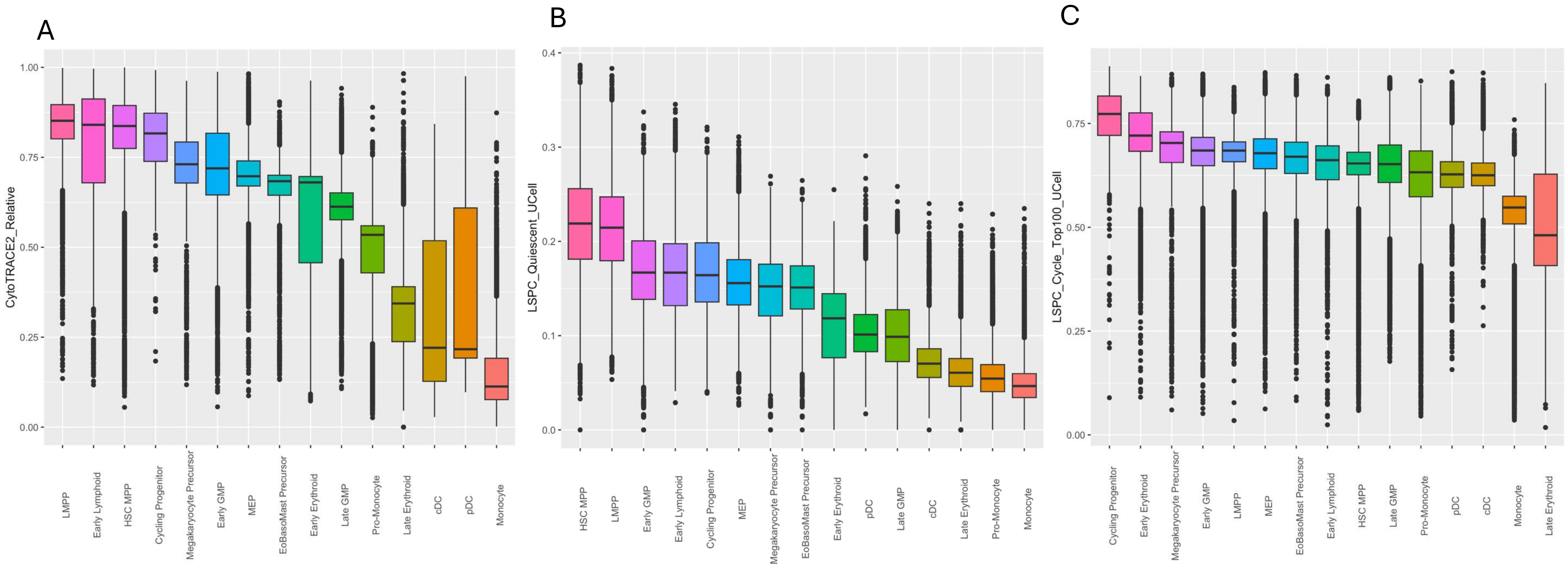

#### Supplementary figure 4. Composition shift at disease progression

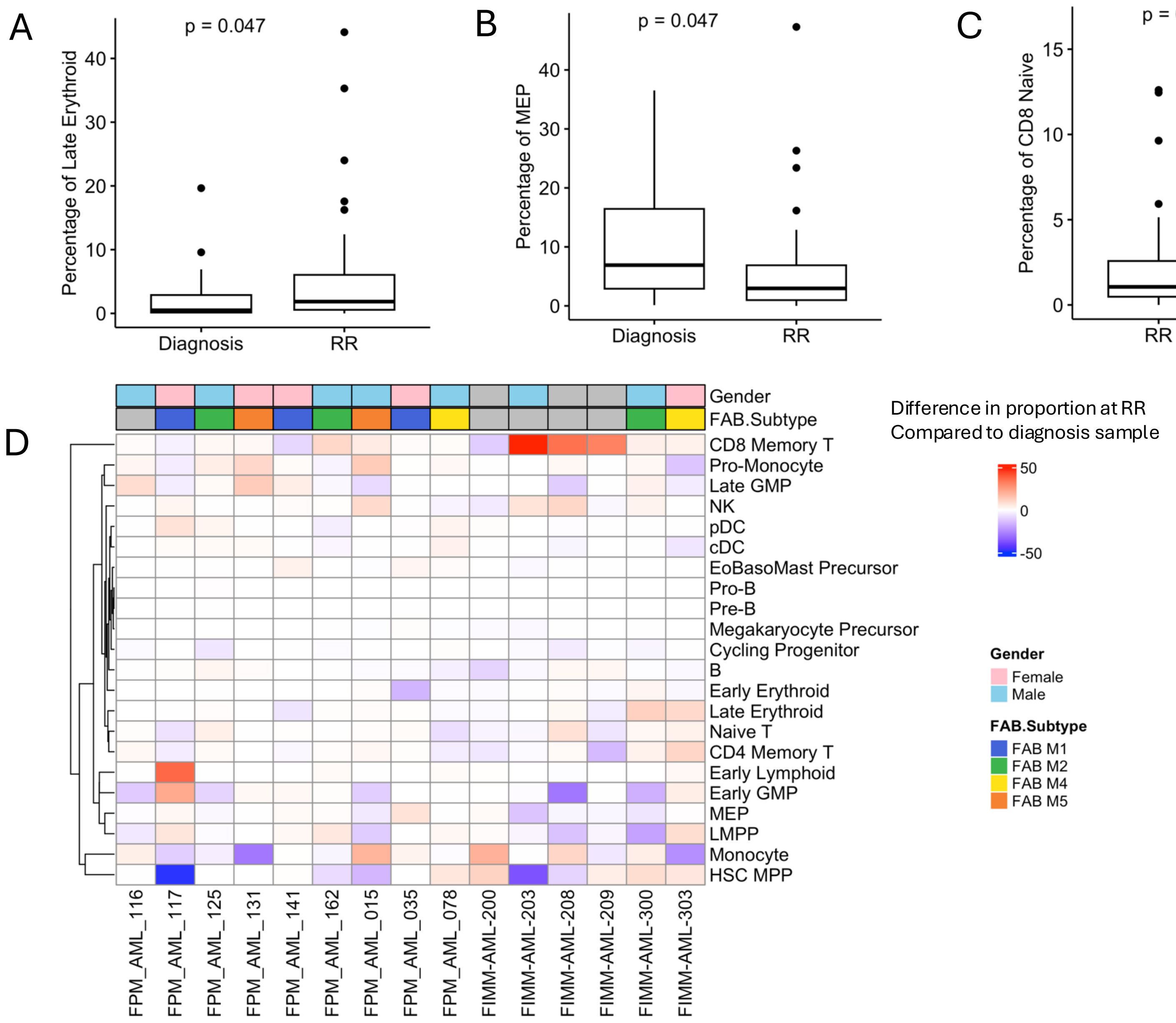

**Supplementary figure 5. Composition shift at disease progression after Cytarabine-based and Venetoclax-based treatment regimen**

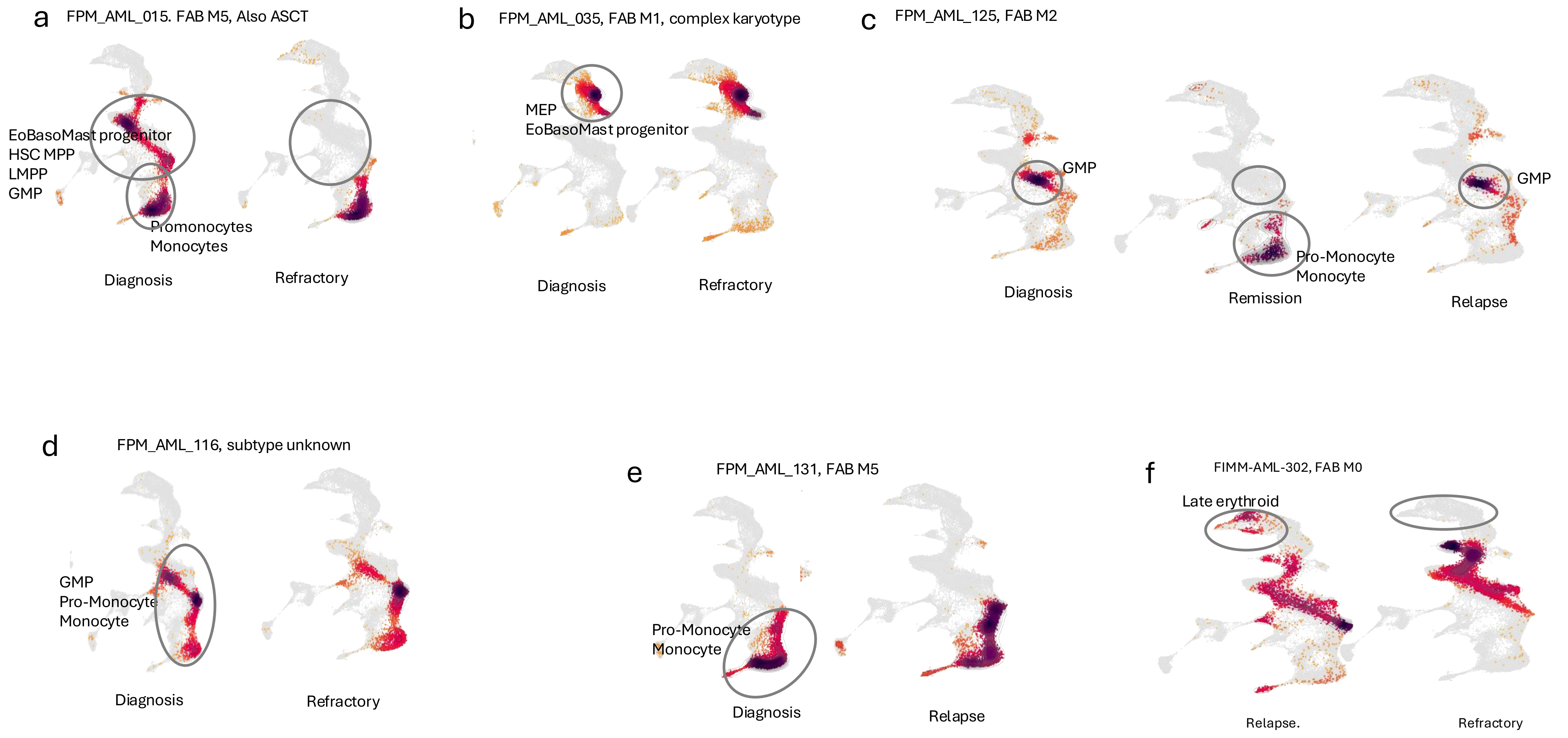

#### Supplementary figure 6. Altered cell-cell communication in AML

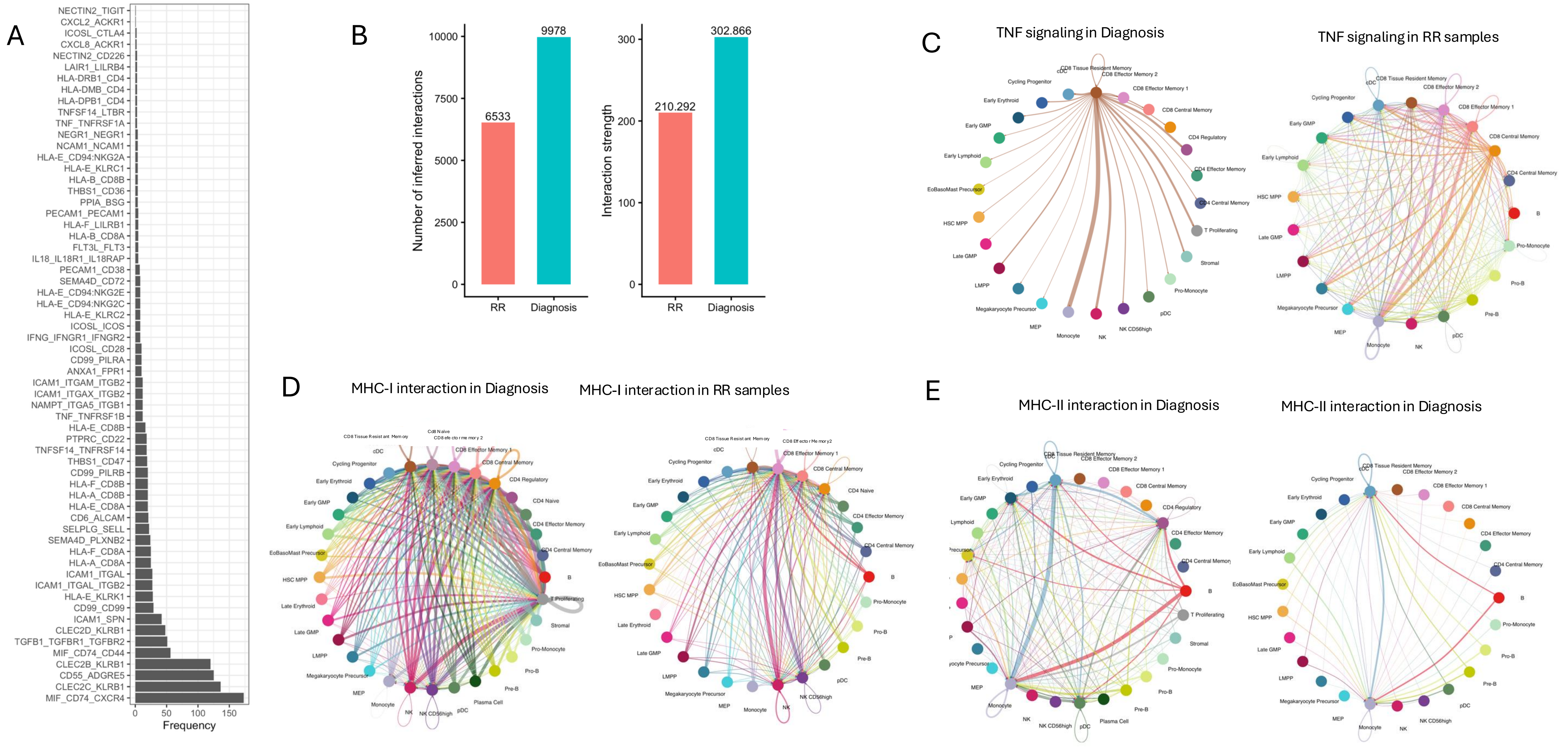

### Supplementary figure 7. Expression of immune related marker genes

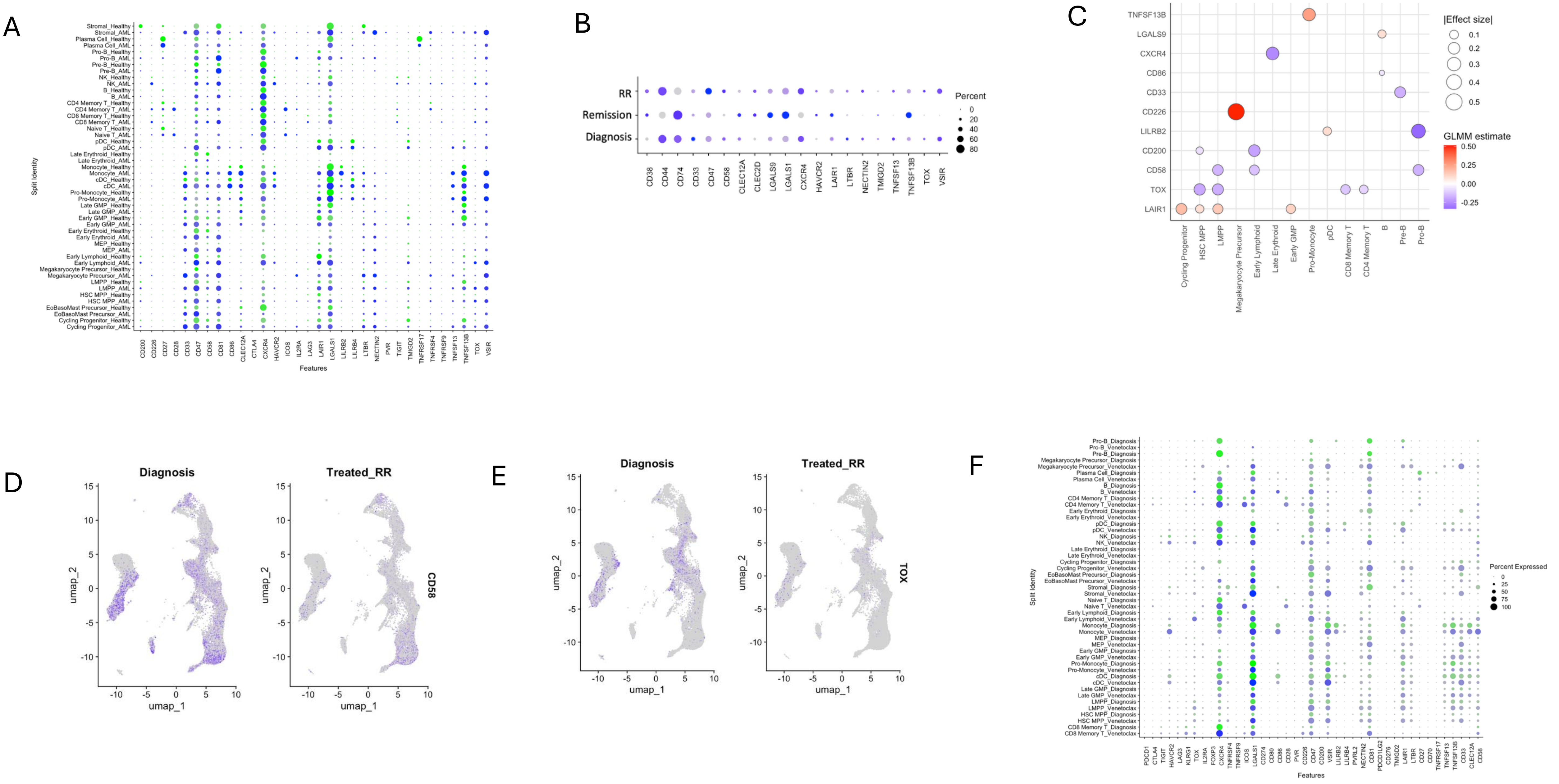

### Supplementary figure 8. Significant cellular composition changes in mutated vs wildtype samples

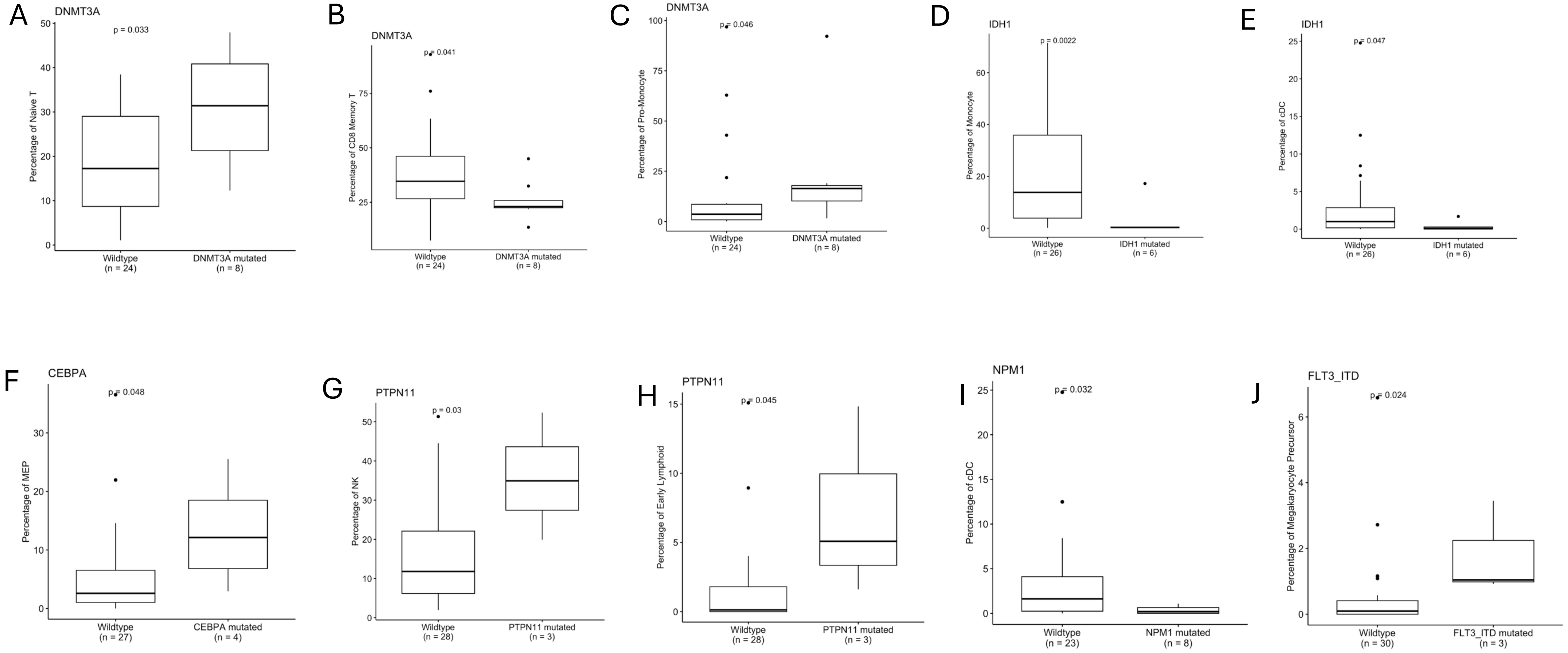

**Supplementary figure captions:**

**Supplementary figure 1. Cellular composition in AML.** A. number of cells for AML cohort  B.Broader cell states. C. Majority of AML samples showed myeloid dominance D. Progenitor clusters showed expression of AML markers D-E. Celltypes with significant differences according to fab subtypes.

**Supplementary figure 2. Composition shift in AML compared to healthy BM samples.** A. Differences in cell composition in AML and Healthy dataset normalized to total cells. B. Differences in cell composition in AML and Healthy dataset normalized to myeloid cells.

**Supplementary figure 3. Functional states in AML myeloid cell populations.** A. Cytotrace score. B. LSPC Quiescent score. C. LSPC Cell Cycle score.

**Supplementary figure 4. Composition shift at disease progression.** A-C. Significant composition shift at RR compared to diagnosis samples. D. Longitudinal samples show patient specific cellular changes

**Supplementary figure 5. Composition shift at disease progression after Cytarabine-based and Venetoclax-based treatment regimen**
**Supplementary figure 6. Altered cell-cell communication in AML.** A. Counts of interactions in AML enriched communication pathways. B. Number of interactions and interaction strength in diagnosis and RR samples. C-E Esample of pathways present in both diagnosis and RR group, where RR showed differential number and strength of interactions.

**Supplementary figure 7. Expression of immune related marker genes.** A.Significant changes identified by GLMM analysis in AML vs healthy samples.B.Significant changes identified by GLMM analysis in AML disease stage. C. Significant changes identified by GLMM analysis in Cytarabine treated RR samples compared to diagnosis. Red represents higher in RR. D-E- Featureplot showing some immune markers reduced in cytarabine treated RR samples.F. Immune markers in venetoclax treated RR samples compared to diagnosis.

**Supplementary figure 8. Significant cellular composition changes in mutated vs wildtype samples**
